# Proteomic data from leaves of twenty-four sunflower genotypes underwater deficit

**DOI:** 10.1101/2020.06.26.171066

**Authors:** Thierry Balliau, Harold Duruflé, Nicolas Blanchet, Mélisande Blein-Nicolas, Nicolas B. Langlade, Michel Zivy

## Abstract

This article describes how the proteomic data were produced on sunflower plants subjected to water deficit. Twenty-four sunflower genotypes were selected to represent genetic diversity within cultivated sunflower. They included both inbred lines and their hybrids

Water deficit was applied to plants in pots at the vegetative stage using the high-throughput phenotyping platform Heliaphen. Here, we provide proteomic data from sunflower leaves corresponding to the identification of 3062 proteins and the quantification of 1211 of them in these 24 genotypes grown in two watering conditions. These data differentiate both treatment and the different genotypes and constitute a valuable resource to the community to study adaptation of crops to drought and the molecular basis of heterosis.

## Value of the Data

- Drought is an important issue for crop yield that will get worse in the context of climate change. Sunflower, the fourth oilseed crop in the world, is particularly impacted by this stress.
- Heterosis is a phenomenon commonly used to improve yield. To analyse its impact on the response to environmental constraints and particularly to drought, twenty-four genotypes of cultivated sunflower comprising four maintainer lines, four restorer lines and their 16 corresponding hybrids were subjected to two treatments (Well-Watered or Water-Deficit).
- Plants were managed on the outdoor Heliaphen high-throughput phenotyping platform. Leaf samples were collected at the end of the treatments for proteomic analysis.
- This dataset provides identification data for 3062 proteins of sunflower leaves and quantification data for 1211 of them in 24 genotypes grown in two conditions.
- It provides unique access to the genetic variability of the sunflower’s molecular response to water deficit.

## 1. Data

Climate change is a current issue of major concern because of its potential effects on biodiversity and the agricultural sector. Better understanding the adaptation of plants to this recent phenomenon is therefore a major interest for crop science and society. *Helianthus annuus* L., the domesticated sunflower, is the fourth most important oilseed crop in the world [3] and is promising for agriculture adaptation because it can maintain stable yields across a wide variety of environmental conditions, especially during drought stress [1]. It constitutes an archetypical systems biology model as drought stress response involves many molecular pathways and subsequent physiological processes.

In this data article, we are sharing the proteomic data of 24 genotypes of sunflower grown in two environmental conditions in the outdoor Heliaphen platform. These datasets are part of a larger project that integrates other omics data at different biological levels (like ecophysiological data described in [2]) and could be associated to diverse studies.

The raw data associated with this article (data from the mass spectrometer in mzXML format as well as annotated spectra and quantitative data can be found at the following link https://doi.org/10.15454/TW59-P718 or directly at http://moulon.inra.fr/protic/sunrise. Protein identification data are provided as a supplemental file and can also be found at http://moulon.inra.fr/protic/sunrise. Parameters used for mass spectrometry analyses are shown in Table 1.

**Table 1:** Parameters used for mass spectrometry analyses.

| XParameters |  |
| --- | --- |
| Identification Software | X! tandem Piledriver (2015.04.01.1) |
|  | http://www.thegpm.org/ |
| Filtering and inference software | X!TandemPipeline 3.4.3 "Elastine Durcie" |
| Filters | Proteins: log (e-value) < -5 |
|  | Proteins: minimum 2 peptides |
|  | Peptides : e-value < 0.01 |
| Digestion | Trypsine |
|  | No semi-tryptic peptide allowed |
|  | 1 misscleavage allowed in first pass and 5 in refine pass |
| Modifications | **Fixed** |
|  | Carbamidomethylation of Cys residues = +57.04 |
|  | **Possible** |
|  | Oxidation of Met and Trp residues = +15.99 |
|  | N-ter acetylation = +42.01 |
|  | N-ter deamidation of Gln residues = -17.02 |
|  | Nter deamidation on carbamidomethylated Cys= -17,02 |
|  | Loss of H<sub>2</sub>O on N-ter Glu residues = -18.01 |
|  | deamidation of Gln and Asn residues = +0.98 |
|  | N-ter Met excision |

## 2. Experimental Design, plant material and growth conditions

The experiment was performed from May to July 2013 in the outdoor Heliaphen phenotyping platform at the Institut National de la Recherche Agronomique (INRA) station, Auzeville, France (43°31’41.8”N, 1°29’58.6”E) as previously described in [4]. Bleach-sterilized seeds were germinated on Petri dishes with Apron XL and Celeste solutions (Syngenta, Basel, Switzerland) for 78 h at 28°C. Germinated plantlets were transplanted in individual pots filled with 15 L of P.A.M.2 potting soil (Proveen distributed by Soprimex, Chateaurenard, Bouches-du-Rhône, France) and covered with a 3-mm-thick polystyrene sheet to prevent soil evaporation. After 17 days after germination (DAG), plants were fertilized with 500ml of Peter’s Professional 17-07-27 (0,6g/l) and extra mix composed of oligo-element Hortilon (0,46g/l) solution. At 21 DAG, Polyaxe at 5mg/l was applied on foliage against thrips.

In total, 144 plants, corresponding to 24 genotypes (4 males and 4 females and their 16 hybrids obtained by crossing) were grown in two (Well-Watered; WW or Water-Deficit; WD) conditions done in triplicate. Each pot was adequately fertilized and irrigated as in [5] before the beginning of the water deficit application. Pots were saturated with water 35 DAG and after excessive water was drained (~ for two hours), pots were weighed to obtain the full soil water retention mass. Then, irrigation was stopped at 38 DAG (~20-leaf stage corresponding to bud formation phase (stage R1 or R3; [6]) for WD plants (as in work by [4]). All pots were covered with a 3 mm layer of polystyrene sheets at the collar level to limit soil evaporation. Soil evaporation was estimated according to [7]. Both WW and WD plants were weighed three or four times per day by the Heliaphen robot to estimate transpiration [4]. WW plants were rewatered at each weighing by the robot to maintain soil water at the retention capacity. Pairs of WD and WW plants were harvested when the FTSW of the stressed plant reached 0.1 (occurring between the 42 and the 47 DAG). Two out of three SF342 plants under control condition died and we could not produce data for those.

At harvest, leaves for molecular analysis were cuts without their petiole and immediately frozen in liquid nitrogen from 11 h to 13 h. On sunflower, the mature leaf developmental stage corresponds to a dark green leaf, assumed to be experiencing its highest photosynthetic rate and having recently reached its maximum size [8]. More precisely, we defined the mature leaf as positioned at three-fifths (0.60 ± 0.04 SD) of the plant (leaf 20 ± 2.3 SD) [2]. The selected leaf harvested for the molecular analysis was the n+1 leaves of the mature.

## 3. Proteomic

### 3.1. Protein extraction

Leaf proteins were extracted using the TCA-acetone protocol described in [9]. Protein digestion was performed according to the liquid digestion protocol described in [10]. Briefly, proteins of the TCA-acetone pellet were solubilized in a buffer containing urea, thiourea, dithiothreitol, Tris-Hcl pH 8.8 and a zwitterionic acid labile surfactant (ZALSI). Proteins were alkylated by using iodoacetamide and digestion by trypsine was performed after dilution in an ammonium bicarbonate solution. Digestion was stopped by adding trifluoroacetic acid, that also allowed ZALS cleavage. The resulting peptide mix was desalted by using C18 solid solid phase extraction. Eluted peptides were then speedvac dried.

LC-MS/MS was performed as described in [10]. Peptides (400 ng) were solubilized in a solution containing 2% acetonitrile and 0.1% formic acid. LC-MS/MS was performed by using an Eksigent nanoLC Ultra 2D nanoHPLC (SCIEX) coupled to a Qexactive mass spectrometer (Thermo, Waltham, MA, USA). After desalting on a trapcolumn, peptides were submitted to a gradient of 5 to 35% ACN that was performed in 40 min. Data-dependent MS analysis was performed with a full scan was at a 75,000 resolution and MS/MS scan at a 17,500 resolution. The isolation window was set to 3 m/z. MS/MS was repeated for the 8 most intense ions detected in full scan and dynamic exclusion was set to 40 s.

### 3.2. Identification of proteins by LC-MS/MS

Protein identification was performed by searching the Heliagen database using genome HanXRQv1 [1] with the X!Tandem search engine [12]. Data filtering and protein inference were performed by using X!TandemPipeline 3.3.4 [13]. Trypsin digestion was declared with respectively one and five possible miss cleavages in the first and refine pass respectively. Only proteins identified with at least two different peptides in the same sample were considered. [14]

### 3.3. Bioinformatics annotation of proteins and quantification

Quantification was operated using the MassChroQ software [14]. Only proteins quantified with at least 2 specific peptides that were present in at least 90% of the samples were selected for analysis. Functional annotation of proteins is given according to the INRAE Sunflower Bioinformatics Resources (www.heliagene.org/HanXRQ-SUNRISE/). All the LC-MS/MS data have been deposited at PROTICdb (https://doi.org/10.15454/TW59-P718).

## Acknowledgments

These data were produced with the funding of the French National Research Agency (ANR SUNRISE ANR-11-BTBR-0005). This work was part of the “Laboratoire d’Excellence (LABEX)” TULIP (ANR-10-LABX-41).

## Notes

http://moulon.inra.fr/protic/sunrise

### Competing Interest Statement

The authors have declared no competing interest.

## References

[1] H. Badouin, J. Gouzy, C.J. Grassa, F. Murat, S.E. Staton, L. Cottret, C. Lelandais-Brière, G.L. Owens, S. Carrère, B. Mayjonade, L. Legrand, N. Gill, N.C. Kane, J.E. Bowers, S. Hubner, A. Bellec, A. Bérard, H. Bergès, N. Blanchet, M.C. Boniface, D. Brunel, O. Catrice, N. Chaidir, C. Claudel, C. Donnadieu, T. Faraut, G. Fievet, N. Helmstetter, M. King, S.J. Knapp, Z. Lai, M.C. Le Paslier, Y. Lippi, L. Lorenzon, J.R. Mandel, G. Marage, G. Marchand, E. Marquand, E. Bret-Mestries, E. Morien, S. Nambeesan, T. Nguyen, P. Pegot-Espagnet, N. Pouilly, F. Raftis, E. Sallet, T. Schiex, J. Thomas, C. Vandecasteele, D. Varès, F. Vear, S. Vautrin, M. Crespi, B. Mangin, J.M. Burke, J. Salse, S. Muños, P. Vincourt, L.H. Rieseberg, N.B. Langlade, The sunflower genome provides insights into oil metabolism, flowering and Asterid evolution, Nature. 546 (2017) 148–152. doi:10.1038/nature22380.

[2] N. Blanchet, P. Casadebaig, P. Debaeke, H. Duruflé, L. Gody, F. Gosseau, N.B. Langlade, P. Maury, Data describing the eco-physiological responses of twenty-four sunflower genotypes to water deficit, Data Br. (2018). doi:10.1016/J.DIB.2018.10.045.

[3] F. USDA, Oilseeds: World Markets and Trade, 2019. https://www.fas.usda.gov/data/oilseeds-world-markets-and-trade (accessed October 1, 2019).

[4] F. Gosseau, N. Blanchet, D. Varès, P. Burger, D. Campergue, C. Colombet, L. Gody, J.F. Liévin, B. Mangin, G. Tison, P. Vincourt, P. Casadebaig, N. Langlade, Heliaphen, an outdoor high-throughput phenotyping platform for genetic studies and crop modeling, Front. Plant Sci. 9 (2019) 1908. doi:10.3389/fpls.2018.01908.

[5] D. Rengel, S. Arribat, P. Maury, M.-L. Martin-Magniette, T. Hourlier, M. Laporte, D. Varès, S. Carrère, P. Grieu, S. Balzergue, J. Gouzy, P. Vincourt, N.B. Langlade, A Gene-Phenotype Network Based on Genetic Variability for Drought Responses Reveals Key Physiological Processes in Controlled and Natural Environments, PLoS One. 7 (2012) e45249. doi:10.1371/journal.pone.0045249.

[6] A.A. Schneiter, J.F. Miller, Description of Sunflower Growth Stages1, 1981. doi:10.2135/cropsci1981.0011183X002100060024x.

[7] G. Marchand, B. Mayjonade, D. Varès, N. Blanchet, M.-C. Boniface, P. Maury, F.N. Andrianasolo, P. Burger, P. Debaeke, P. Casadebaig, P. Vincourt, N.B. Langlade, A biomarker based on gene expression indicates plant water status in controlled and natural environments, Plant. Cell Environ. 36 (2013) 2175–2189. doi:10.1111/pce.12127.

[8] F.N. Andrianasolo, P. Casadebaig, N. Langlade, P. Debaeke, P. Maury, Effects of plant growth stage and leaf againg on the response of transpiration and photosynthesis to water deficit in sunflower, Funct. Plant Biol. 43 (2016) 797–805. doi:10.1016/j.neurobiolaging.2008.03.012.

[9] V. Méchin, C. Damerval, M. Zivy, Total protein extraction with TCA-acetone., Methods Mol. Biol. 355 (2007) 1–8. doi:10.1385/1-59745-227-0:1.

[10] V. Hervé, H. Duruflé, H. San Clemente, C. Albenne, T. Balliau, M. Zivy, C. Dunand, E. Jamet, An enlarged cell wall proteome of Arabidopsis thaliana rosettes, Proteomics. 16 (2016). doi:10.1002/pmic.201600290.

[11] T. Balliau, M. Blein-Nicolas, M. Zivy, Evaluation of Optimized Tube-Gel Methods of Sample Preparation for Large-Scale Plant Proteomics, Proteomes. 6 (2018) 6. doi:10.3390/proteomes6010006.

[12] R. Craig, J.P. Cortens, R.C. Beavis, Open source system for analyzing, validating, and storing protein identification data, J. Proteome Res. 3 (2004) 1234–1242. doi:10.1021/pr049882h.

[13] O. Langella, B. Valot, T. Balliau, M. Blein-Nicolas, L. Bonhomme, M. Zivy, X! TandemPipeline: A Tool to Manage Sequence Redundancy for Protein Inference and Phosphosite Identification, J Proteome Res. 16 (2017) 494–503. doi:10.1021/acs.jproteome.6b00632.

[14] B. Valot, O. Langella, E. Nano, M. Zivy, MassChroQ: a versatile tool for mass spectrometry quantification, Proteomics. 11 (2011) 3572–3577. doi:10.1002/pmic.201100120.

